# *GsERF* enhances aluminum tolerance through an ethylene-mediated pathway in *Arabidopsis thaliana*

**DOI:** 10.1101/2020.07.01.182253

**Authors:** Lu Li, Xingang Li, Ce Yang, Yanbo Cheng, Zhandong Cai, Hai Nian, Qibin Ma

**Author notes:** These authors contributed equally to this work (Lu Li, Xingang Li). Correspondence: College of Agriculture, South China Agricultural University.

## Abstract

The ethylene response factor (ERF) transcription factor is a subfamily of AP2/ERF superfamily in plants, which plays multiple roles in plant growth and development as well as stress response. In this study, we found that the *GsERF* gene from BW69 line of wild soybean held a constitutive expression pattern and induced by aluminum stress with more transcripts in soybean root. The putative GsERF protein containing an AP2 domain was in the nucleus and transactivation activity. In addition, the overexpression of the *GsERF* gene enhanced root relative length rate in Arabidopsis and shallow staining by hematoxylin under the treatments of AlCl_3_. The ethylene synthesis related genes such as *ACS4, ACS5* and *ACS6* are upregulated in the *GsERF* overexpressed plants than those in wild type plants under the treatment of AlCl_3_. Furthermore, expression levels of stress/ABA-responsive marker genes, including *ABI1, ABI2, ABI4, ABI5, RD29B* and *RD22* in transgenic lines compared with those in wild type Arabidopsis were affected by AlCl_3_ treatments. Taken together, the results indicate that overexpression of *GsERF* may enhance aluminum tolerance through an ethylene-mediated pathway and/or ABA signaling pathway in *Arabidopsis thaliana*.

## INTRODUCTION

The toxicity of aluminum (Al) is a major limiting factor for crop production in acidic soils, which account for about 50% of the world’s arable land (Uexküll *et al*., 1995). When the pH value of soil is lower than 5.0, aluminum will exist in the form of ions, and Al^3+^ will dissolve into the soil which will strongly inhibit root growth and function, thus reduce crop yield (Kochian, 1995). Within a species, plant species and varieties vary widely in their ability to tolerate aluminum toxicity. Some plant species or varieties have evolved high levels of tolerance mechanisms in order to survive in acidic soils. Wild soybean in South China has been growing in acid soil for a long time and there is no lack of tolerance resources, which plays an important role in improving the stress resistance of soybean (Zeng *et al*., 2012).

ERF transcription factor (ethylene response factor) is a subfamily of AP2/ERF superfamily, which can be divided into three categories according to the number of AP2/ERF domains: AP2, ERF and RAV (Zhang *et al*., 2008). The ERF family of proteins contains an AP2/ERF domain consisting of a highly conserved 58-60 amino acids that binds to multiple cis-acting elements, including GCC box and DRE/CRT, etc (Ohme-Takagi *et al*., 1995; Liu *et al*., 1998). This domain is the main functional region of ERF family genes (Riechmann *et al*., 1998). It is found that ethylene response factors (ERF) not only play an important role in plant growth and development, but also play a very important role in plant response to stress (Riechmann *et al*., 1998). Previous studies have shown that ERF family genes are involved in plant growth and development in rice, Arabidopsis and other plants. Such as OsERF1 is constitutively expressed in different organs of rice, and is upregulated by ethylene, overexpression of *OsERF1* in *Arabidopsis thaliana* promotes the expression of ethylene responsive gene *PDF1.2* and b-chitinase, and significantly affects the growth and development of transgenic Arabidopsis (Hu *et al*., 2008). AtERF71/HRE2 can activate the expression of downstream genes by binding with GCC box and DRE/CRT, regulate the expansion of root cells and play an important role in root development (Lee *et al*., 2015). Julien Pirrello found that overexpression of *Sl-ERF2* gene in transgenic tomato lines can lead to early seed germination and enhanced hook formation in dark growth seedlings. Recently, the transcription factor ERF139 was found in poplar to regulate the expansion of xylem cells and the deposition of secondary cell walls (Wessels *et al*., 2019).

In recent years, more and more ERF family genes are found to function in stress tolerance in plants. Under drought stress, it was found that overexpressing rice genes *OsERF71, OsERF101* and *OsERF48* can enhance drought resistance of rice (Jin *et al*., 2018; Jung *et al*., 2017; Li *et al*., 2018). Heterologous overexpression of soybean gene *GmERF3* can enhance tobacco drought resistance (Zhang *et al*., 2009). Overexpressed *AtERF019* can enhance drought resistance in Arabidopsis (Scarpeci *et al*., 2016). Under salt stress, allogeneic overexpression of *GmERF135* in Arabidopsis can enhance the salt tolerance of Arabidopsis plants. Meanwhile, *GmERF135* can promote the growth of transgenic hairy roots under salt stress (Zhao *et al*., 2019). In wheat, overexpression of ERF1-V can enhance the salt tolerance of wheat, and heterologous overexpression of *GmERF7* can enhance the salt tolerance of tobacco (Zhai *et al*., 2013; Xing *et al*., 2017). Under alkaline stress conditions, heterologous overexpression in Arabidopsis thaliana, *GsERF71* and *GsERF6* from wild soybeans, and *VaERF3* from red beans, could enhance the resistance of Arabidopsis to alkali stress (Li *et al*., 2020; Yu *et al*., 2017; Yu *et al*., 2016). Overexpression of ZmEREB180 in maize can enhance maize flood tolerance (Yu *et al*., 2019). Heterologous overexpression of *VaERF092* and *ERF105* enhanced Arabidopsis cold tolerance (Bolt *et al*., 2017; Sun *et al*., 2019). Overexpression of *GmERF75* in Arabidopsis can enhance osmotic stress tolerance of Arabidopsis, and GmERF75 can promote osmotic stress tolerance in transgenic hairy roots (Zhao *et al*., 2019). In addition, ERF genes can also enhance plant resistance to pathogens. *AtERF14* was found in regulating plant defense response (Oñate-Sánchez *et al*., 2007). In Arabidopsis, *ERF11* and *ERF15* can positively regulate immunity to *Pseudomonas syringae* (Zhang, 2015; Zheng *et al*., 2019). In soybean, *GmERF13* and *GmERF5* can enhance the resistance to *Phytophthora sojae* (Zhao *et al*., 2017; Dong *et al*., 2015).

However, no ERF family gene has been reported to be involved in the response of plants to acid-tolerant aluminum stress. In this study, qRT-PCR analysis showed that aluminum could rapidly induce the expression of *GsERF* gene in wild soybean, and *GsERF* gene showed a constitutive expression pattern in wild soybean, which was expressed in soybean leaves more than that in root, but the expression level of *GsERF* gene in soybean root tip was the highest under the condition of aluminum stress. This discovery prompted us to further explore the response mechanism of *GsERF* gene to aluminum stress.

## RESULTS

### Isolation and sequence analysis of *GsERF* gene

In this study, the full-length cDNA sequence of *GsERF* gene was cloned from the BW69 line of *Glycine soja* which is tolerant to Al toxicity. The primers were designed according to the *GsERF* homologue gene in *Glycine max, Glyma09g52900*. The *GsERF* gene contains a complete open reading frame (ORF) of 369 bp which is 99 % identical to *Glyma09g52900* based on the genome database from Phytozome, and encodes a protein of 123 amino acids. The predicted GsERF protein contained a conserved DNA-binding domain (AP2/ERF domain) of 58 amino acids which is reported to be the primary functional area. Alignment analysis revealed that GsERF was 68% to 97% similar to other homologous genes in domain ratio (Fig.1). The analysis of ERF gene family indicated that GsERF is a member of B-2 subgroup members (Zhang *et al*., 2008). A number of ERF family genes have been reported to have related functions, many of them can play a role in the plant in the face of biotic and abiotic stress. Through phylogenetic tree analysis, we found that GsERF and GmERF5 are closely related in the same branch (Fig.2).

**Figure 1.**
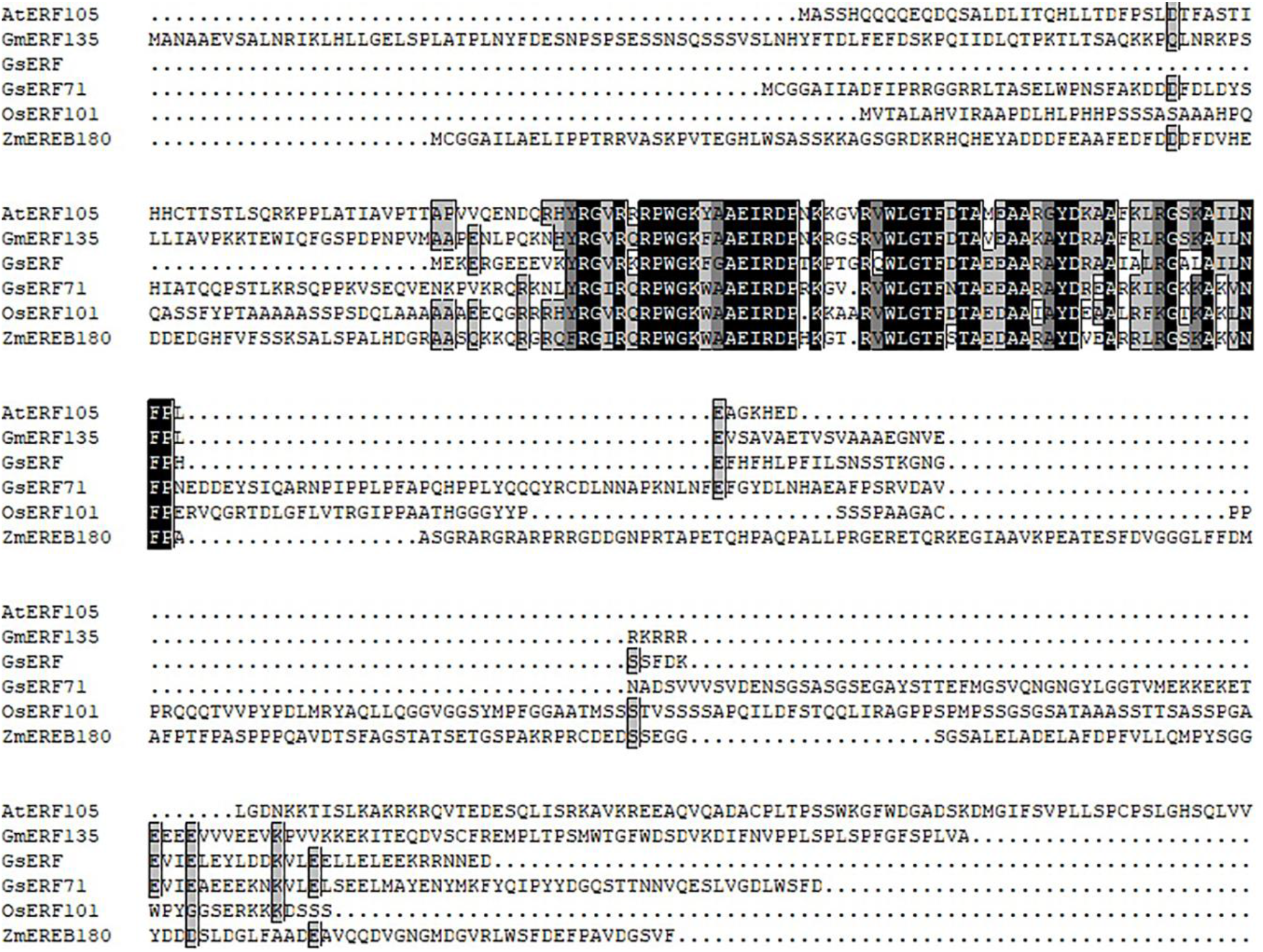
Sequence alignment of AP2 domain by DNAMAN. The shaded part of the figure is the AP2 domain. The protein sequences of the selected ERF genes were obtained from Phytozome or Genebank, the Accession Numbers were shown follows: AtERF105 (NP_568755.1), GmERF135 (Glyma.17G145300), OsERF71 (XP_015643752.1), OsERF101 (Os04g32620), ZmEREB180 (NC_024459.2).

**Figure 2.**
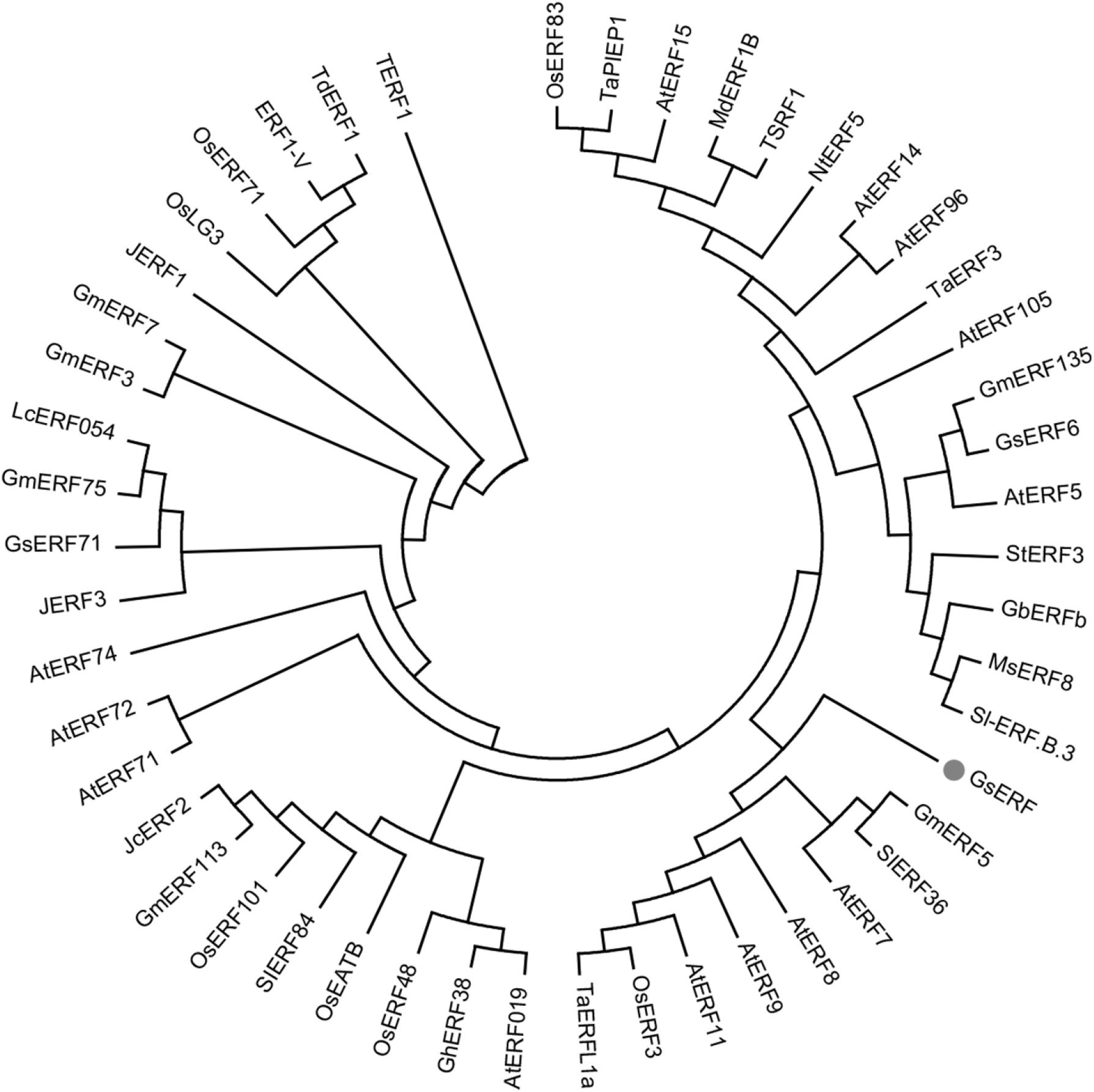
Phylogenetic relationship was constructed with 48 transcription factors of the ERF family associated with stress resistance. The protein sequences of the selected ERF genes were obtained from Phytozome or Genebank, the Accession Numbers were shown in the supplementary material.

### *GsERF* expression pattern analysis

Quantitative real-time PCR (qRT-PCR) was performed to assess the transcript levels of *GsERF* in BW69 plants. The qRT-PCR results showed that *GsERF* was a constitutive expression pattern rich in roots, stems and leaves. Under the condition of aluminum stress, the expression level of *GsERF* in roots, stems and leaves was significantly increased, especially at the root tip of 1cm with the expression levels of 13 times (Fig.3C). Under the treatments of different aluminum concentrations time courses, the results showed that *GsERF* could be rapidly induced by aluminum stress, and the transcripts of *GsERF* reached a maximum level at 4 h, and then the mRNA transcripts of *GsERF* begin to decline (Fig.3A). Under the treatments of different aluminum concentrations, the *GsERF* transcripts increased with AlCl3 concentrations, and when the concentration of AlCl3 was 100 μM, the level of *GsERF* mRNA was 25 times that of the control (Fig.3B).

**Figure 3.**
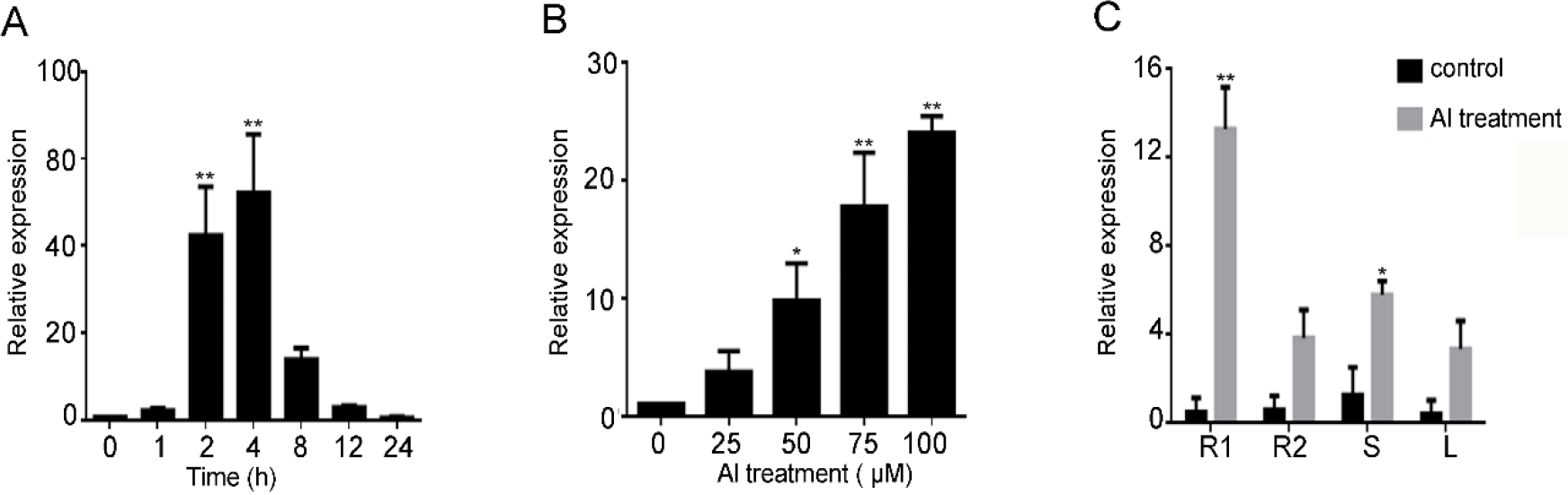
Expression patterns of *GsERF* were detected in different tissues and under AlCl_3_ treatments. A, *GsERF* expression in roots exposed to 30 μM AlCl_3_ for 0 to 24 h. B, *GsERF* expression in roots exposed to 0 to 100 μM AlCl_3_ for 6 h. C, *GsERF* expression in soybean root apices (R1,0-1cm), root apices (R2,0=-2cm),stems (S) and leaves (L) in the absence or presence of Al stress. Values are means ± SD (n = 3). Asterisks show significant differences between control and Al treatments using student’s t-test: *, P<0.05; **, P<0.01.

### GsERF is a nuclear protein with transactivation activity

To determine the cellular localization of GsERF protein, its localization was analysed by expressing a gene encoding a GsERF–eGFP fusion protein under the control of the CaMV35S promoter in tobacco epidermal cells. The empty vector (PCAMBIA1302-eGFP) was used as the control. As shown in Fig.4A, the eGFP protein was distributed in the whole cells, while the GsERF1-eGFP fusion protein was only visualized in the cell nuclei. The results clearly indicated that GmERF5 is a nuclear-localized protein.

**Figure 4.**
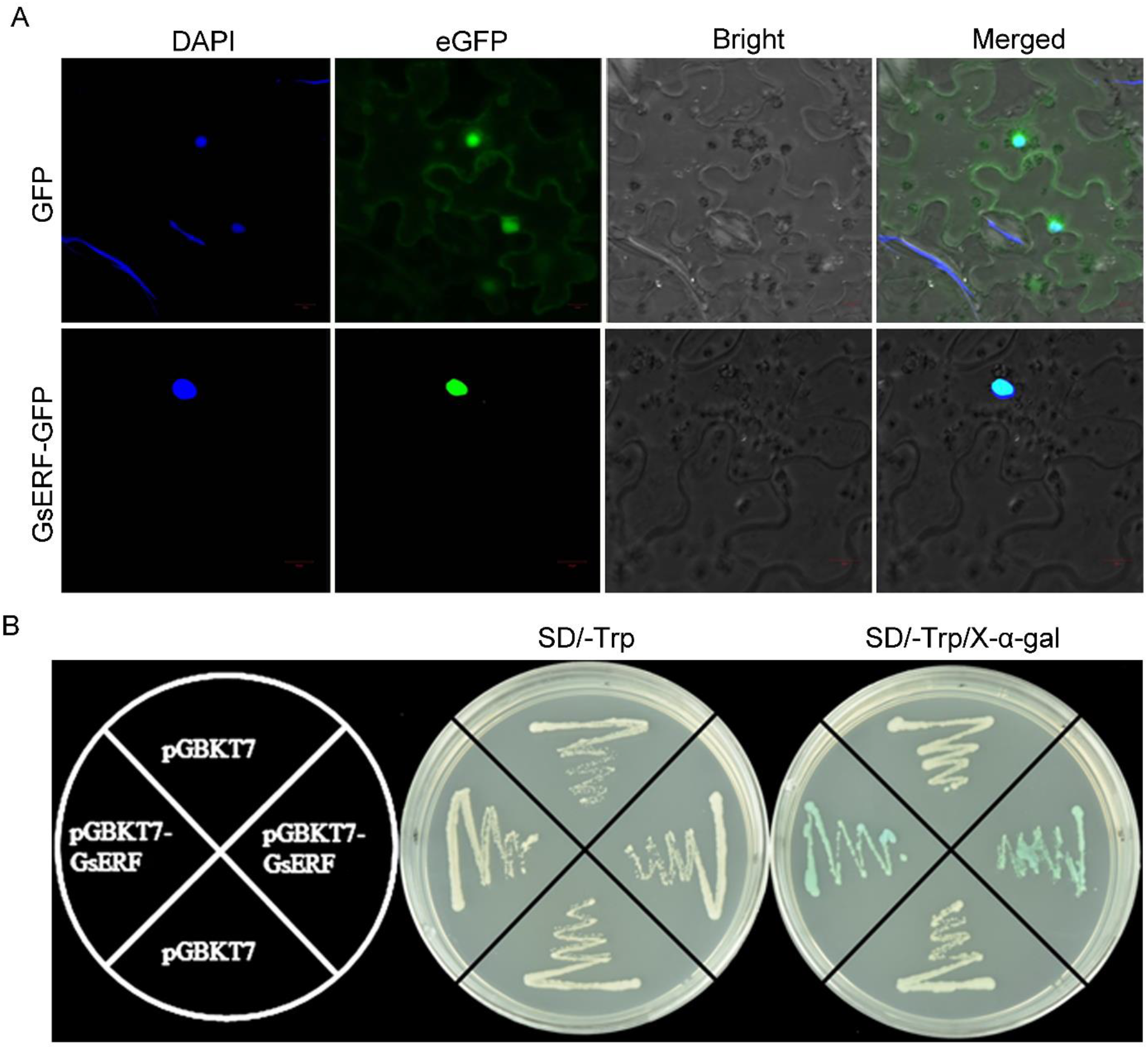
GsERF protein is localized in the nucleus and possesses transactivation activity. A, Nuclear localization of the GsERF protein in leaf epidermis cells of Nicotiana benthamiana. Nicotiana benthamiana leaves transiently expressing GFP alone (upper) and GsERF-GFP (bottom) proteins were observed with a confocal microscope (厂家, 生产国家). B, Transactivation assay of the truncated GsERF proteins. Full-length of GsERF were fused with the GAL4 DNA-binding domain and then expressed in yeast strain Y2H gold. The transformed yeast cells were plated and grown on control plates (SD/-Trp) or selective plates (SD/-Trp +X-α-gal).

To determine whether GsERF could act as a transcriptional activator, the yeast two-hybrid analysis was used. Full-length of *GsERF* gene were fused with the GAL4 DNA-binding domain and then expressed in yeast strain Y2H gold to identify the transcriptional activation activity by growing the yeast cells on control plates (SD/-Trp) or selective plates (SD/-Trp +X-α-gal). Yeast cells containing only the pGBKT7 plasmid of GAL4 DNA binding domain were used as the negative control. The results showed that only yeast colonies carrying GsERF could activate the expression of reporter gene and make the colonies blue on the selective plate (Fig.4B).

### Overexpression of *GsERF* enhanced plant Al tolerance

To investigate the effect of *GsERF* under aluminum stress, GsERF was overexpressed in Arabidopsis to obtain the transgenic lines (Fig.S1). Then, three homozygous lines with high expression were selected for the phenotype identification. Under the treatment of AlCl3, the growth of *GsERF* transgenic plants and WT plants were significantly inhibited, but root growth of *GsERF* overexpression (OX) lines was less inhibited than that of WT plants. The statistical results also showed that the relative root elongation of *GsERF* transgenic plants was significantly higher than that of WT plants (Fig.5A&B). The proline content of transgenic plants and wild-type plants increased after aluminum treatment, but the proline content in *GsERF* transgenic plants was much higher than that of wild-type plants (Fig.5C).

**Figure 5.**
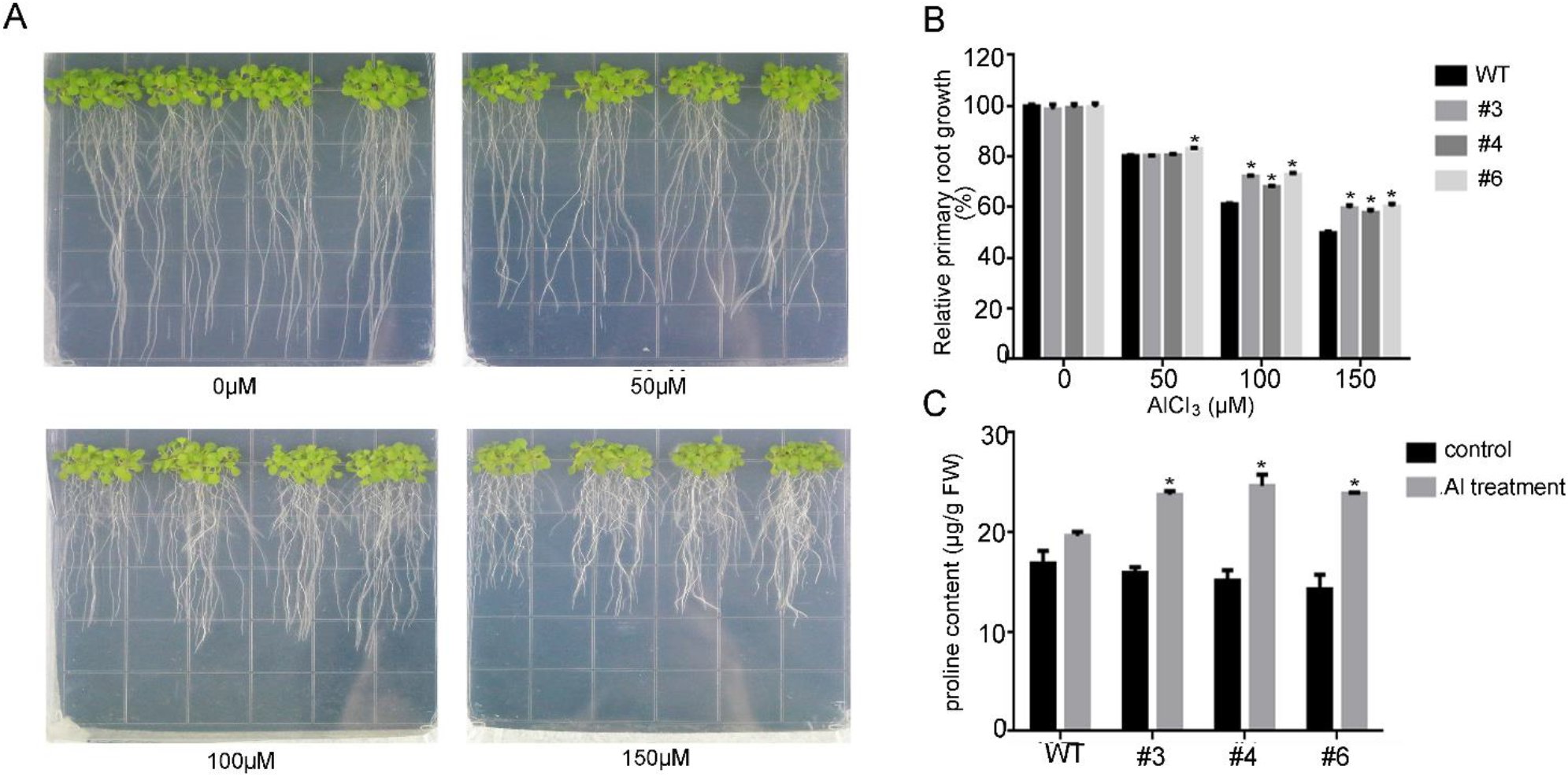
GsERF enhanced the resistance of Arabidopsis plants to Al stress. A, The root growth of wild type and GsERF overexpression Arabidopsis with or without Al treatment. B, The relative root length growth was calculated. C, The free proline contents in wild type and GsERF overexpression Arabidopsis. Data are mean values ± SD (*P<0.05; Student’s t-test). All experiments described earlier were carried out with three biological repetitions.

In order to verify the role of *GsERF* in soybean, we obtained overexpressed and RNAi interfering hair root lines through soybean hairy root transformation. We detected that GsERF mRNA levels were 38 times higher in overexpressing lines than those of WT. However, *GsERF* mRNA levels were 80 percent lower in RNAi lines than that of WT (Fig.6A). To detect the dying of hairy roots, the hairy roots of Arabidopsis were stained with hematoxylin when they were treated in the solution containing 50 μM AlCl3 for 6 hours. The staining results showed that the hairy roots in control were stronger than those in OX-GsERF transgenic lines, and weaker than those in RNAi-GsERF transgenic lines (Fig.6B). The results indicated that the binding amount of Al^3+^ in hairy roots is the least in OX-GsERF lines, while the binding amount of Al^3+^ in hairy roots is the most in RNAi-GsERF lines. Taken together, these results suggested that *GsERF* overexpression can enhance the tolerance of Arabidopsis and soybean to aluminum stress.

**Figure 6.**
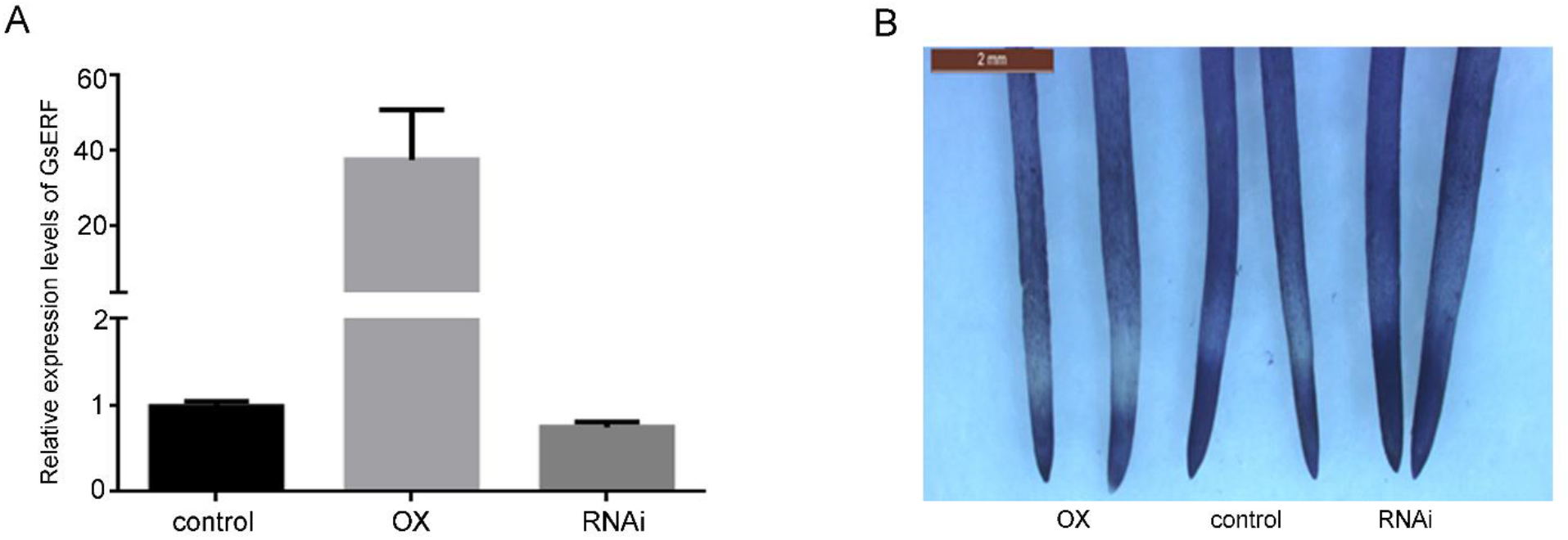
The root tip staining phenotype of hairy roots. A, the expression level of *GsERF* in the control, OX-GsERF and RNAi-GsERF hairy roots. B, the control, OX-GsERF and RNAi-GsERF hairy roots were stained with hematoxylin dye after Al treatment. The control, OX-GsERF and RNAi-GsERF hairy roots were treated with 50 μM AlCl_3_ for 12 h. Data are mean values ± SD, all experiments described earlier were carried out with three biological repetitions.

### *GsERF* enhances aluminum tolerance through an ethylene-mediated pathway

To understand the molecular mechanism of *GsERF* tolerant to Al stress, some genes from ERF family members were carried out to investigate their responses to Al stress and ACC (ethylene precursor) content was determined in Arabidopsis. On the base of *GsERF* expression up-regulated by ethylene (ET) (Fig. S2), we examined the changes of ACC (ethylene precursor) content in *GsERF* overexpression (OX) lines and wild type (WT) in Arabidopsis after aluminum treatment 10 days. We found that the content of ACC in *GsERF* overexpression plants after aluminum treatment was higher than that of wild-type plants, while there was a little difference in ACC content between wild type plants and overexpressed plants without aluminum treatment (Fig.7A). These results suggested that ET signal transduction may be involved in the aluminum-tolerant pathway induced by *GsERF* gene. To further confirm this hypothesis, we analyzed the transcription levels of genes associated with ET synthesis related genes. the qRT-PCR results showed that the expression levels of *ACS4, ACS5* and *ACS6* genes related to ethylene synthesis was significantly increased in the *GsERF* overexpressed plants than those in wild type plants under the treatment of AlCl3 (Fig. 7B, C&D).

**Figure 7.**
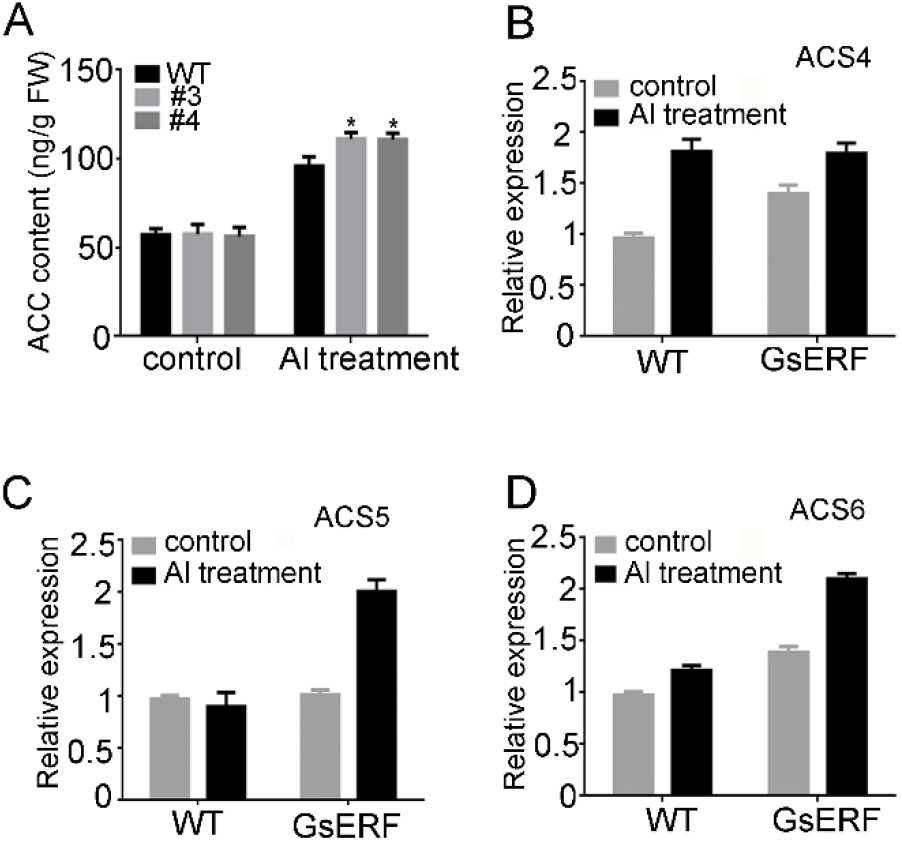
ACC content and the expression of ACC biosynthesis genes in GsERF-overexpressing and wild type Arabidopsis. A, ACC contents in GsERF-overexpressing and wild type Arabidopsis under control and Al treatment conditions. B-D, Real-time PCR analysis of the expression of ACC biosynthesis genes under control and Al treatment conditions. Data are mean values ± SD (*P<0.05; Student’s t-test). Error bars represent the standard error of three replicates. WT: wild type. #3 and #4: transgenic lines of GsERF in T3 generation.

## DISCUSSION

Aluminum toxicity has a great influence on the root of crops, which will directly affect the yield of crops. Therefore, it is important in theory and practice to find new aluminum tolerance genes and determine their functions. AP2/ERF family is one of the largest families of plant transcription factors, whose members are involved in many aspects of plant development and respond to many environmental stresses (Klay *et al*., 2018). In this study, *GsERF* gene encoding an ERF transcription factor was isolated from gene expression profile resistant to aluminum stress using wild soybean BW69 line (Zeng *et al*., 2012). Sequence analysis showed that GsERF protein has a highly conserved AP2 domain with typical characteristic of the b-3 subgroup of the ERF superfamily (Fig.1) (Zhang *et al*., 2008). Furthermore, GsERF protein localizes in the nucleus and has a characteristic of self-activation activity like many other ERF transcription factors (Fig.4).We speculated that may GsERF protein as a transcription factor hold the potential functions of ERF family members.

According to previous studies, members of the AP2/ERF family have important functions in plants facing various environments, different growth and development stages. For example, members of the B-3 subgroup, where *OsERF48* can enhance the tolerance of plants to drought and salt stresses (Jung *et al*., 2017), *AtERF15* can enhance the positive regulation of the immune response in Arabidopsis (Zhang, 2015), and *AtERF096* can reduce the water loss rate in Arabidopsis (Wang *et al*., 2015), NtERF5 enhances resistance to tobacco Mosaic virus, and *GsERF6* regulates plant tolerance to bicarbonate in Arabidopsis (Yu *et al*., 2016). In addition, the ERF genes also play a key role in plant growth and development. For example, *MdERF1B* can regulate the biosynthesis of apple anthocyanin and procyanidins (Zhang *et al*., 2018), and *OsERF1* significantly affects the growth and development of transgenic Arabidopsis (Hu *et al*., 2008). However, no ERF family genes have been reported to be involved in the response to Al stress in plants. In present study, *GsERF* with a constitutive expression pattern was rapidly induced by aluminum stress with richest transcripts under AlCl3 treatment (Fig.3). This result suggested *GsERF* may play certain role in aluminum stress.

In order to verify whether the gene GsERF has the function of acid-tolerant aluminum, we induced soybean hairy root transformation and heterologous transformation of Arabidopsis thaliana, combining the colorimetric properties of hematoxylin, hematoxylin and bluish-purple when complexed with Al, therefore, visual evaluation of stained roots can be used to detect the accumulation of Al in root tissues. In recent years, there have been more and more reports using soybean hairy root transformation to verify the function of soybean aluminum tolerance genes. Among them, GmME1-OE hairy root tip hematoxylin staining is lighter and the aluminum content is lower, indicating that aluminum resistance is stronger. The hairy roots overexpressing GmGRPL had obvious aluminum resistance than the control. Under aluminum stress, the aluminum content in the hairy roots overexpressing GmGRPL was lower than that of the control, and the hairy roots overexpressing GmGRPL had higher antioxidant capacity. In our study, we found that after culturing in the presence of aluminum, after hematoxylin staining, the hairy roots of the overexpressed lines were lighter than the root tips of the control lines, and the control lines were stained more than the interference lines. Light, which means that the GsERF overexpression hairy root tip aluminum content is lower than the control, and the color of the root tip Al complexed with hematoxylin is lighter. Previous studies have shown that root tip elongation is inhibited when plants are subjected to aluminum stress, so root tip elongation is one of the indicators of aluminum tolerance (Wang *et al*., 2006). Similarly, in GsERF transgenic Arabidopsis thaliana, we found that the relative root elongation of transgenic lines was significantly greater than that of the wild type in the presence of aluminum, and the proline content of transgenic plants was higher than that of the wild type after aluminum treatment. Known research results indicate that the plant’s proline content will increase under stress, and a large amount of accumulated proline will help the plant reduce the damage caused by stress. Combined with our experimental results, GsERF gene can enhance the tolerance of transgenic Arabidopsis to aluminum stress. However, in our study, we found that when GsERF overexpressing plants were subjected to aluminum stress, they produced more ethylene than wild-type plants, and the ethylene synthesis related gene ASC4, ASC5 and ASC6 expression levels increased. The results were similar to those found in other studies, which found that ASC1, ASC2, and ASC5 genes associated with ethylene synthesis increased in NaHCO3 stress in GmERF7 overexpressed lines (Yu *et al*., 2016). In addition, protein phosphatase 2A can reduce the toxicity of cadmium by regulating ethylene production in Arabidopsis, among which ASC2 and ASC6 are found to be up-regulated under cadmium stress (Chen *et al*., 2020). According to previous studies, plant hormones are involved in the response to stress. When plants are under stress, various hormones in the body will react, and different hormones may interact to form a network of mutual exchanges to resist external pressure (Ku *et al*., 2018). In this study, the ethylene synthesis related genes such as *ASC4, ASC5* and *ASC6* were induced with much higher levels by *GsERF* under aluminum stress (Fig.7). In addition, abscisic acid content were measured to investigate its potential role in regulating Al stress. The determine results showed that there was a little difference on the ABA content between *GsERF* overexpressing lines and wild type in Arabidopsis plants (Fig.S3). However, some genes involved in the abscisic acid signaling pathway were found to have changed expression levels in *GsERF* overexpressing lines (Fig.8). Under aluminum stress, the transcripts of *ABI1* and *ABI2* in *GsERF*-overexpressing lines were significantly lower than those in wild type (Fig.8). Studies have shown that *ABI1* and *ABI2* play a key role in abscisic acid signal transduction and are negative regulators in abscisic acid signal transduction (Seo *et al*., 2010). While the transcripts of *ABI4* and *ABI5* in *GsERF*-overexpressing lines was higher than those of WT, which are positive regulators of abscisic acid signal transduction (Finkelstein *et al*., 2000; Finkelstein *et al*., 1998). Furthermore, *RD29B* is significantly up-regulated in *GsERF* overexpressed lines (Fig.8). It is well known that *RD29B* is mainly involved in drought, salt stress and abscisic acid response through independent abscisic acid pathways resulting in higher permeability and stress tolerance of plants (Msanne *et al*., 2011; Shinozaki, 1994). The results suggested that GsERF gene may regulate plants tolerant to aluminum stress through ET pathway and/or the interaction between ethylene and abscisic acid.

**Figure 8.**
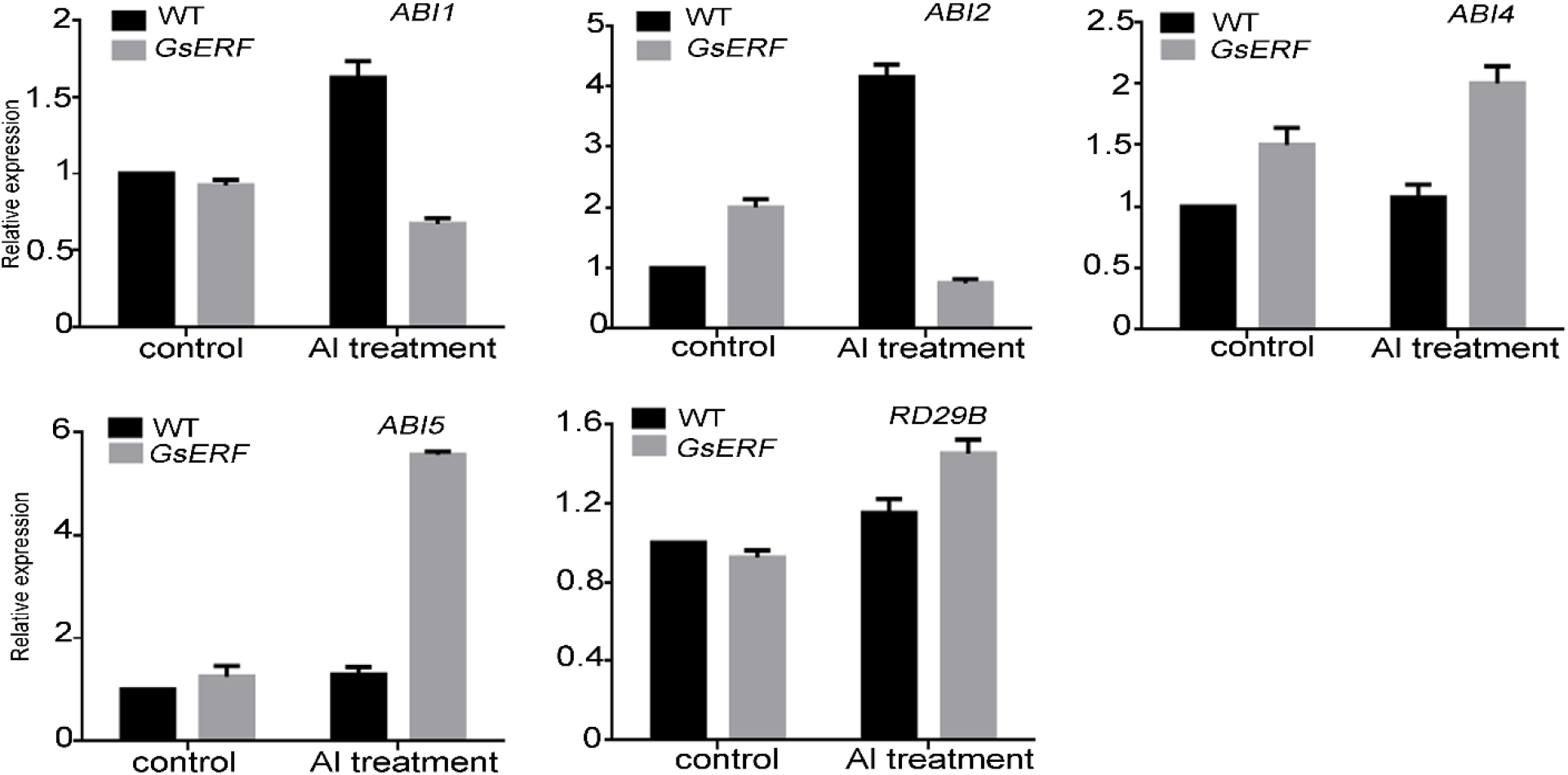
Expression levels of ABA transport-related genes in GsERF-overexpressing and wild type Arabidopsis. Seedlings with approximately 1-cm roots were grown in agar medium containing 0 or 150 mM AlCl_3_ for 10 days. Error bars indicate standard error of the means (SD) based on three technical replicates. Data are mean values ± SD.

## MATERIALS and METHODS

### Plant material and stress treatment

The wild soybean seeds from BW69 line were grown in a growth chamber maintained at 28°C /25°C and 70% relative humidity with a 14 h light / 10 h dark cycles. The seeds germinated in vermiculite, then seedlings of uniform growth were selected and cultured in nutrient solution (pH 5.8) for three days. The solution was renewed daily. After 3 days, the seedlings were transplanted into 0.5 mM CaCl2 (pH 4.5) solution were used for aluminum treatments. For ethylene stress, the hydroponic seedlings were placed in an airtight plexiglass chamber, and ethylene gas was released after 2 ml 40% ethephon and 1 g NaHCO_3_ were dissolved in 200 ml H_2_O (Zhang *et al*., 2009).

The Arabidopsis ecotype Columbia (Col-0) seeds were germinated and grown in the growth chamber with following conditions: 22-24°C, 60% relative humidity, 100 mol photons m^-2^s^-1^ 16 h light and 8 h dark cycles. Seeds are planted on demand on vegetative soil, from germination to harvest. For the analysis of gene expression of Arabidopsis thaliana, seeds were planted on 1/2 MS agar plates in darkness for 4 days at 4 °C, and then placed in the growth chamber. Then, seedlings in root length 1cm were to 1/2 MS solid medium in the absence or presence of AlCl_3_. After 10 days, the whole plants were collected as samples, and each sample contained at least 10 plants.

### RNA isolation, cDNA synthesis and quantitative real-time PCR

Total RNA was isolated using TRIzol (Tiangen Biotech, Beijing, China) and cDNA syntheses were performed by PrimeScript RT reagent kit (Takara). The qRT-PCR analyses were performed using SYBR Premix ExTaq TM II Mix (TaKaRa, Shiga, Japan). *Actin3* (GenBank accession no.) was used as reference in wild soybean. The Arabidopsis housekeeping gene *actin* (GenBank accession no.) was used as reference in Arabidopsis. The data were analyzed with the 2^-ΔΔCT^ method (Livak *et al*., 2001). The primers used for qRT PCR were listed in Supplementary Table S1.

### GsERF gene isolation and sequence analysis

The *GsERF* gene was isolated from the wild soybean BW69 line. The full sequence of *GsERF* was amplified by PCR primer pairs 5’ - GGATCACGCCTCAAGTT −3’ and 5’-CGAACCCTAAATCATCAG −3’. The PCR products were inserted into the multiple cloning site of the pLB vector (Tiangen Biotech, Beijing, China) and sent for sequencing. Multiple alignments of sequences analysis were performed using DNAMAN software. Homology analysis of *GsERF* and the other 44 reference ERF superfamily genes were performed using MAGE6.0 software by a neighbor-joining method. The amino acid sequences were obtained from GenBank (http://www.ncbi.nlm.nih.gov/genbank/) or Phytozome (http://phytozome.jgi.doe.gov/pz/portal.html).

### Subcellular localization analysis

To analyze subcellular localization of GsERF protein, the full-length of *GsERF* was inserted into the *NcoI/SpeI* site of pCAMBIA1302 vector to generate GsERF-eGFP construct. The fusion construct pCAMBIA1302-GsERF-eGFP was transformed into tobacco epidermal cells. After 2-3 days, the green fluorescence signals in tobacco epidermal cells were observed under the confocal laser-scanning microscope (SP5, Leica, Wetzlar,Germany)(Sparkes *et al*., 2006).

### *In vitro* transcriptional activation assay

For the transactivation assay, the full-length of *GsERF* was inserted into the *Eco*RI/*Bam*HI sites of pGBKT7 vector. The constructs of pGBKT7-GsERF were transformed into yeast strain Y2Hgold and grown on SD/-Trp medium at 30 °C for 3 days. After selection of the yeast transformants carrying the GsERF gene on SD (-Trp) medium, they were transferred to SD (-Trp, X-α-Gal) medium to identify the transcriptional activation. The empty plasmid were used as negative control.

### Arabidopsis transformation and soybean hairy root transformation

The Arabidopsis ecotype Col-0 was used for transformation. With CaMV 35S as the promotor, the full coding region of *GsERF* was inserted into the plant expression binary vector pTF101.1 to generate pTF101.1-GsERF. The constructs were transformed to Agrobaterium tumefaciens strain GV3101 and then the targeted gene was transferred into Arabidopsis plants by floral dip method (Clough, 2010).

Five-day-old seedlings with unfolded cotyledons were used for soybean hairy root production. For the RNAi construct, 233 bp of the *GsERF* coding region was cloned and inserted into the pMU103 vector. The overexpression vector and RNAi interference vector were transferred into A. rhizogenes strain K599 and then transformed by hypocotyl injection (Guo *et al*., 2011). The empty plant expression binary vector pTF101.1 were used as control.

### Hematoxylin staining

The expression of *GsERF* gene in hairy root lines was analyzed, and appropriate hairy root lines were selected for subsequent experiments. Hairy roots were used for 0, 25 μM AlCl3 treatments (0.5 mM CaCl2, pH 4.5) for 6 h. After AlCl3 treatment, the hairy roots were washed three times with sterilized water and were stained with hematoxylin dye. The dyed roots were washed in sterile water for half an hour and then observed and photographed by Leica S8APO stereo microscope (Leica, Germany) (Rincón *et al*., 1992).

### Phenotype analysis of Arabidopsis tolerant to Acidic aluminum stress

To analyze the phenotypes of *GsERF* overexpression (OX) and wild type (WT) Arabidopsis under aluminum stress, the seeds of T3 GsERF overexpression and WT were used in this study. The seed surface was sterilized with 10% sodium hypochlorite for 10 minutes and subsequently washed in deionized water. Then, the sterilized seeds were grown on 1/2 MS agar plates in darkness for 4 days at 4 °C. Then the plate is cultivated upright, 22-24°C, 60% relative humidity, 100 mol photons m-2s-1 16 h light and 8 h dark cycles. Then seedlings with root length of 1 cm were selected and transferred to 1/2 MS agar medium (pH 4.5) with different AlCl3 concentrations. After 10 days, measure the main root length and take pictures with a camera.

### Physiological indices assay

GsERF overexpression and WT lines were treated with or without aluminum for 10 days, and the whole plant was selected as a sample. The free proline content was measured as the method described in detail previously (Zhang *et al*., 2009). Ethylene precursor (ACC) and abscisic acid content was determined using an enzyme-linked immunosorbent assay (ELISA)(Yang *et al*., 2001).

### Statistical analysis

All experiments with each group were performed at least in triplicate. Data were reported as mean ± SD. All data were analysed by Graphpad Prism 6.01 software by the t test to assess significant differences among means.

## ACKNOWLEDGMENTS

This work was supported by grants from the Major Project of New Varieties Cultivation of Genetically Modified Organisms (2016ZX08004002-007), the National Natural Science Foundation of China (31771816, 31971965), the Special Supervision on Quality and Safety of Agricultural Products of the Ministry of Agriculture and Rural Areas (4100-C17106, 21301091702101); the Key Projects of International Scientific and Technological Innovation Cooperation among Governments under National Key R & D Plan (2018YFE0116900), the China Agricultural Research System (CARS-04-PS09), the Key-Area Research and Development Program of Guangdong Province (2020B020220008) and the Project of Science and Technology of Guangzhou (201804020015)

## AUTHOR CONTRIBUTIONS

Q.M., H.N. and L.L. conceived and designed of the study. L.L., X.L., C.Y., Y.C. and Z.C. conducted the experiment. L.L., X.L., and Q.M. performed data as well as statistical analysis. L.L. wrote the manuscript which was reviewed and edited by X.L., H.N. and Q.M.

**SUPPORTING INFORMATIONS**

